# Synaptic release of acetylcholine rapidly suppresses cortical activity by recruiting muscarinic receptors in layer 4

**DOI:** 10.1101/257980

**Authors:** Rajan Dasgupta, Frederik Seibt, Michael Beierlein

## Abstract

Basal forebrain (BF) cholinergic projections to neocortex dynamically regulate information processing. However, the underlying synaptic and cellular mechanisms remain poorly understood. While synaptically released acetylcholine (ACh) can recruit nicotinic ACh receptors (nAChRs) expressed in distinct types of interneurons, previous work has not defined a clear role for muscarinic ACh receptors (mAChRs) in the fast cholinergic control of cortical activity. To address this question, we employed a slice model of cortical activity and used optogenetics to selectively activate cholinergic afferents. We found that transient ACh increases led to a rapid and persistent suppression of cortical activity, mediated by mAChRs in layer 4 and by nAChRs in layer 2/3. Furthermore, mAChR-dependent cholinergic control was mediated at least in part by a short-latency and long-lasting inhibition of layer 4 excitatory neurons. Thus, the activation of postsynaptic mAChRs is central to the flexible cholinergic control of cortical activity.

## Introduction

Cholinergic afferents from the basal forebrain (BF) to neocortex play critical roles in a diverse set of cognitive functions such as attention (Herrero et al., 2008), learning (Letzkus et al., 2011) and sensory processing (Fu et al., 2014). Activation of neuromodulatory systems, including cholinergic and adrenergic projections, can induce changes in internal cortical state, defined by a switch from spontaneous, low frequency, rhythmic activity to desynchronized activity. Such state changes can be global, slow and persistent, as during transitions from sleep to wakefulness (Brown et al., 2012). More recent studies have revealed much more rapid transitions in cortical dynamics *within* the awake state, thereby shaping sensory processing and behavioral performance on a moment-to-moment basis (Crochet and Petersen, 2006; Reimer et al., 2014; McGinley et al., 2015a; Vinck et al., 2015). Neuromodulatory systems are thought to be critically involved in controlling these rapid fluctuations of brain state (Eggermann et al., 2014; Reimer et al., 2016), suggesting a high functional specificity and precision of the underlying circuitry (Muñoz and Rudy, 2014).

BF cholinergic afferents target all cortical layers (Bloem et al., 2014; Wu et al., 2014) and both nAChRs and mAChRs are expressed in a layer-, cell-, and synapse-specific manner (McCormick, 1992; Arroyo et al., 2014; Muñoz and Rudy, 2014; Obermayer et al., 2017). However, the activation of these receptors by synaptically released ACh and the consequences for local circuit dynamics are not well understood. Recent studies have shown that nAChRs expressed in inhibitory interneurons of the superficial cortical layers can mediate rapid feedforward inhibition (Arroyo et al., 2012) or disinhibition (Letzkus et al., 2011; Fu et al., 2014; Letzkus et al., 2015) of cortical circuits. By contrast, activation of mAChRs is thought to generate more gradual and longer-lasting changes in cortical dynamics (Lucas-Meunier et al., 2003; Ballinger et al., 2016). While mAChR agonists can generate a plethora of responses in distinct cell types and synapses (McCormick, 1992; Gil et al., 1997; Kruglikov and Rudy, 2008), only few studies have isolated mAChR-dependent responses induced by endogenous ACh (Hedrick and Waters, 2015). In vivo, BF-evoked changes in cortical activity have been shown to be mediated in part by mAChRs (Goard and Dan, 2009; Eggermann et al., 2014; Muñoz et al., 2017) but the underlying mechanisms are unresolved.

By employing optogenetic activation of cholinergic inputs in a slice model of cortical activity, we demonstrate that ACh release evoked by single pulses rapidly and powerfully inhibited evoked cortical network activity for several seconds. This inhibition was mediated in large part by the activation mAChRs in layer 4, and to a lesser extent by the recruitment of nAChRs in the supragranular layers. We found that synaptically released ACh produced long-lasting mAChR-mediated IPSCs in the majority of layer 4 excitatory neurons and nAChR EPSCs in interneurons of the superficial layers. Our findings reveal that the rapid and reliable recruitment of mAChRs in layer 4 mediates short-latency and persistent control of cortical network activity.

## Results

### Synaptic release of ACh suppresses evoked cortical activity

We investigated the role of cholinergic synaptic signaling in regulating cortical activity by employing optogenetic techniques in somatosensory (barrel) cortical slices of ChAT-ChR2-EYFP mice expressing channelrhodopsin-2 (ChR2) in cholinergic neurons (Zhao et al., 2011). Cortical activity was evoked by applying brief stimulus bursts (4 stimuli, 40 Hz) delivered through extracellular glass electrodes placed in layer 4 and monitored using voltage-clamp recordings from layer 2/3 cells in the same cortical column (Figure 1A). Stimulus bursts generated postsynaptic responses consisting of short-latency monosynaptic EPSCs with little latency jitter as well as long-latency polysynaptic activity (onset: 45.7 ± 6 ms, duration: 678.9 ± 50.9 ms, n = 19 cells) which displayed considerable jitter from trial-to-trial (Figure 1B). Stimulus intensity was adjusted to reliably evoke polysynaptic activity for the majority of trials (90.3 ± 3%, n = 19 cells) in a given recording. As polysynaptic responses are thought to be mediated by recurrent excitatory connections in local cortical networks (Beierlein et al., 2002) we will refer to these responses as recurrent activity. To examine fast cholinergic modulation of recurrent activity, we paired extracellular stimulation in layer 4 with single light pulses (5 ms) centered on the recorded neuron, applied 15 ms prior to the onset of stimulus bursts. This led to a reliable and repeatable suppression of recurrent activity (quantified as the change in EPSC charge transfer, see Materials and Methods) compared to unpaired trials lacking cholinergic stimulation (unpaired: 105 ± 15 pC, paired: 27.8 ± 4 pC, n = 19 cells, p < 0.001, Wilcoxon signed rank test; Figure 1B-E). Similarly, when cells were held in current clamp, optical stimulation led to a reduction of spiking activity (unpaired: 1.18 ± 0.4 spikes per trial, paired: 0.38 ± 0.2 spikes per trial, n = 6 cells, p = 0.01, Wilcoxon signed rank test; Figure 1-figure supplement 1). In contrast, monosynaptic EPSCs evoked by the first two stimuli were unaffected by optical stimulation (unpaired: 197.5 ± 35 pA, paired: 199.6 ± 36 pA, n = 19, p = 0.5, two-tailed paired t-test; Figure 1F) suggesting that under our experimental conditions ACh-mediated modulation of presynaptic glutamate release onto layer 2/3 cells was not prominent. Simultaneous recordings from neighboring layer 2/3 neurons revealed strong covariation of cholinergic suppression of recurrent activity from trial to trial, indicating that cholinergic signaling uniformly suppressed recurrent activity for neurons that form part of the same local network (Figure 1-figure supplement 2). Next, we tested if activity in local inhibitory neuronal networks was similarly reduced by cholinergic signaling by simultaneously recording isolated EPSCs and IPSCs in neighboring neurons, voltage clamped at −70 and 0 mV, respectively (Figure 1G-H). Across cell pairs, suppression of recurrent activity ranged from 71.1% to 1.2% normalized to unpaired trials, with suppression of excitatory and inhibitory activity being virtually identical for a given cell pair (r^2^ = 0.98; Figure 1H), indicating that cholinergic signaling did not alter the ratio of synaptic excitation and inhibition in layer 2/3 during recurrent activity. Taken together, our data indicate that brief activation of cholinergic afferents reliably suppresses recurrent activity in cortical networks.

**Figure 1:**
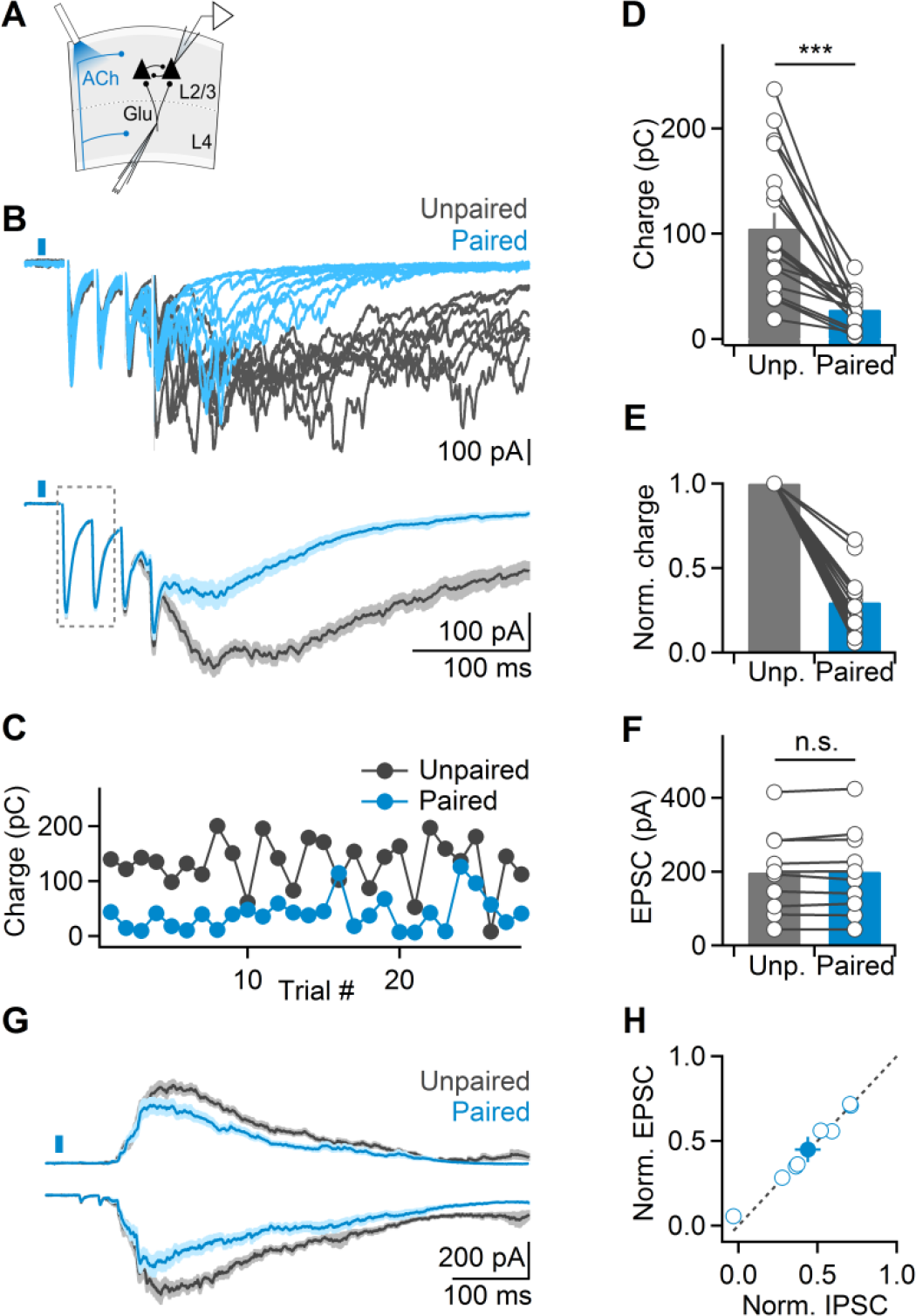
ACh release evoked by single light pulses suppresses evoked cortical recurrent activity. **(A)** Schematic of experimental setup. Cortical recurrent activity was evoked using brief bursts of extracellular stimuli applied in layer 4 and was recorded in voltage clamp in layer 2/3. Cholinergic afferents were activated using single LED pulses (5 ms), 15 ms prior to electrical stimulation. **(B)** *Top:* Representative recording showing multiple trials of recurrent activity, in the absence (unpaired, black traces) or presence of optical stimulation (paired, blue traces). *Bottom:* EPSCs averaged across all unpaired and paired trials. Note lack of amplitude reduction of monosynaptic EPSCs (outlined) **(C)** For the same cell shown in B, plot depicts recurrent activity (quantified as EPSC charge transfer), in paired trials (blue) alternated with unpaired trials (black). **(D)** Summary data showing light-evoked suppression of recurrent activity in layer 2/3 neurons (n = 19 cells). **(E)** Same data as in (D), normalized to unpaired responses. **(F)** Summary data showing average amplitude of monosynaptic EPSC evoked by the first two stimuli (n = 10 cells), for unpaired and paired trials. **(G)** Recurrent activity recorded as EPSCs and IPSCs (black: unpaired, blue: paired) from pairs of neighboring layer 2/3 cells, held at −70 mV and 0 mV, respectively. **(H)** Summary data plotting normalized suppression of EPSCs and IPSCs, for all cell pairs (n = 8). Shaded areas and error bars denote SEM.

### Cholinergic suppression is largely mediated by mAChRs

Both nAChRs and mAChRs are expressed in different types of neocortical neurons (Muñoz and Rudy, 2014), but how these receptors are activated by endogenous ACh to mediate cholinergic control of cortical circuits is not well understood. We found that bath application of atropine to block mAChRs significantly reduced cholinergic suppression (paired: 36.7 ± 5% compared to unpaired trials, atropine: 77.9 ± 4%, n = 10, p < 0.01, Wilcoxon signed rank test; Figure 2A and B), indicating that ACh increases evoked by single stimuli can recruit mAChRs. Application of atropine led to a small increase in recurrent activity in unpaired trials (118.7 ± 9% compared to control, n = 15, p = 0.03, Wilcoxon signed rank test), suggesting that cortical activity is modestly reduced via persistent activation of mAChRs. To determine whether this reduction was due to enhanced levels of ambient ACh in our transgenic mouse model (Kolisnyk et al., 2013), we repeated these experiments in slices derived from wild-type animals. Bath application of atropine still led to an increase in recurrent activity (119 ± 24% compared to control, n = 4, p = 0.23, Wilcoxon signed rank test), indicating that persistent activation of mAChRs was not limited to ChAT-ChR2-EYFP mice.

**Figure 2:**
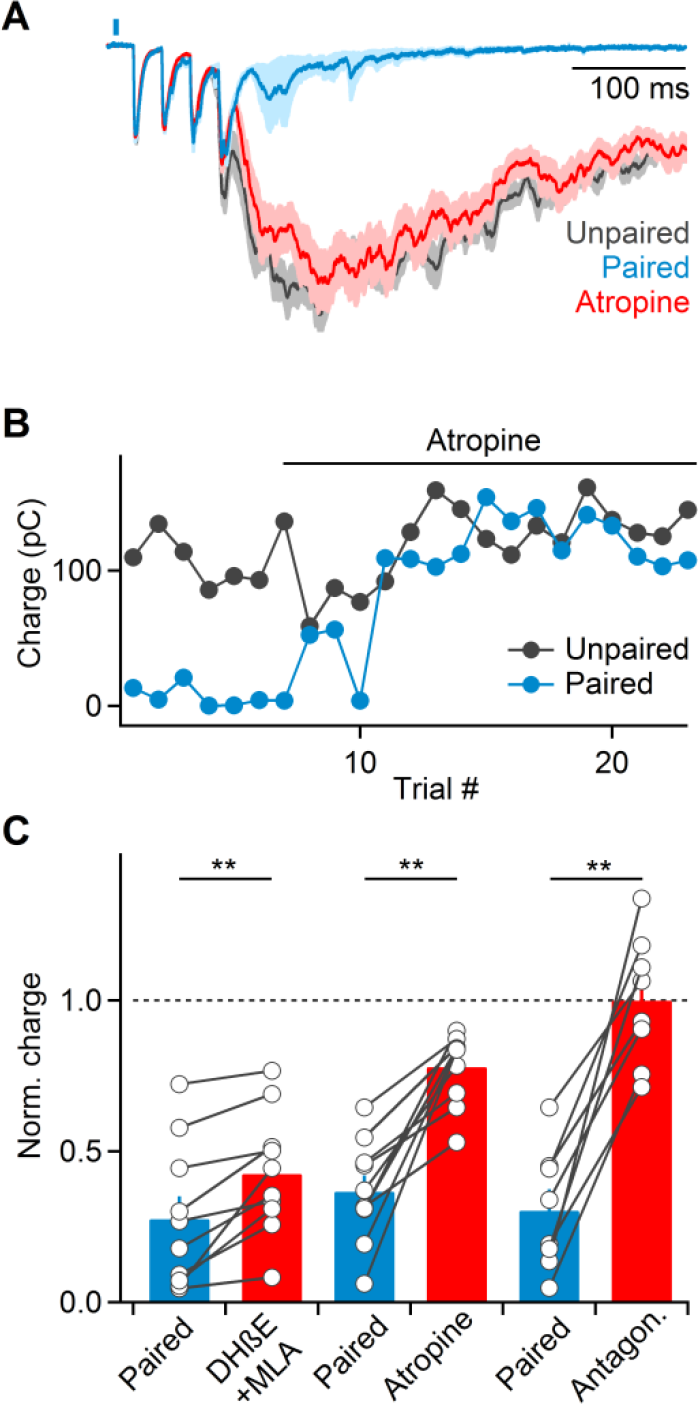
Cholinergic suppression of recurrent activity is mediated by both nAChRs and mAChRs. **(A)** Voltage clamp recordings from a representative layer 2/3 cell showing that bath application of the mAChR antagonist atropine (10 μM) largely blocks cholinergic suppression of recurrent activity (blue: average EPSCs in paired control trials, red: average EPSCs in paired trials following atropine application, gray: average EPSCs in unpaired trials). **(B)** Magnitude of recurrent activity for the same cell across unpaired (black) and paired (blue) during atropine application. **(C)** Summary data of recurrent activity (normalized to activity in unpaired trials) prior to and after bath application of nAChR antagonists (500 nM DHβE + 5 nM MLA, n = 10 cells), atropine (10 μM Atr. n = 10 cells), or both (n = 8 cells). Shaded areas and error bars denote SEM.

Compared to the effects of blocking mAChRs, wash-in of MLA and DHβE to block α7 and non-α7 nAChRs, respectively, led to a smaller but significant reduction of cholinergic suppression (paired: 27.5 ± 7% compared to unpaired trials; MLA and DHβE: 42.5 ± 6%, n = 10, p < 0.01, Wilcoxon signed rank test; Figure 2C). Furthermore, MLA and DHβE application did not lead to an increase in recurrent activity in unpaired trials (96.4 ± 7% compared to control, n = 7, p = 0.25, Wilcoxon signed rank test) suggesting that tonic activation of nAChRs is not prominent.

It is possible that optical stimuli triggered the release of GABA from BF afferents (Saunders et al., 2015) and/or from local ChAT-positive GABAergic neurons expressing ChR2 (von Engelhardt et al., 2007), resulting in suppression of recurrent activity. However, we found that the combined application of both mAChR and nAChR antagonists completely eliminated suppression of recurrent activity (control: 30.4 ± 7% compared to unpaired trials; antagonists: 99.9 ± 8%, n = 8, p < 0.01, Wilcoxon signed rank test; Figure 2C), indicating that light-evoked effects on recurrent activity were largely mediated by ACh.

### Activation of mAChRs leads to prolonged suppression of recurrent activity

The crucial role of mAChRs in the suppression of recurrent activity predicts that suppression should be long-lasting. To examine this possibility, we progressively increased the delay between optical activation of cholinergic afferents and extracellular stimulation to evoke recurrent activity. Suppression of recurrent activity was maximal for delays of 1 and 2 seconds and decreased for 5 second delays, with delays of 8 seconds no longer yielding significant reductions in activity (Figure 3A and B).

**Figure 3:**
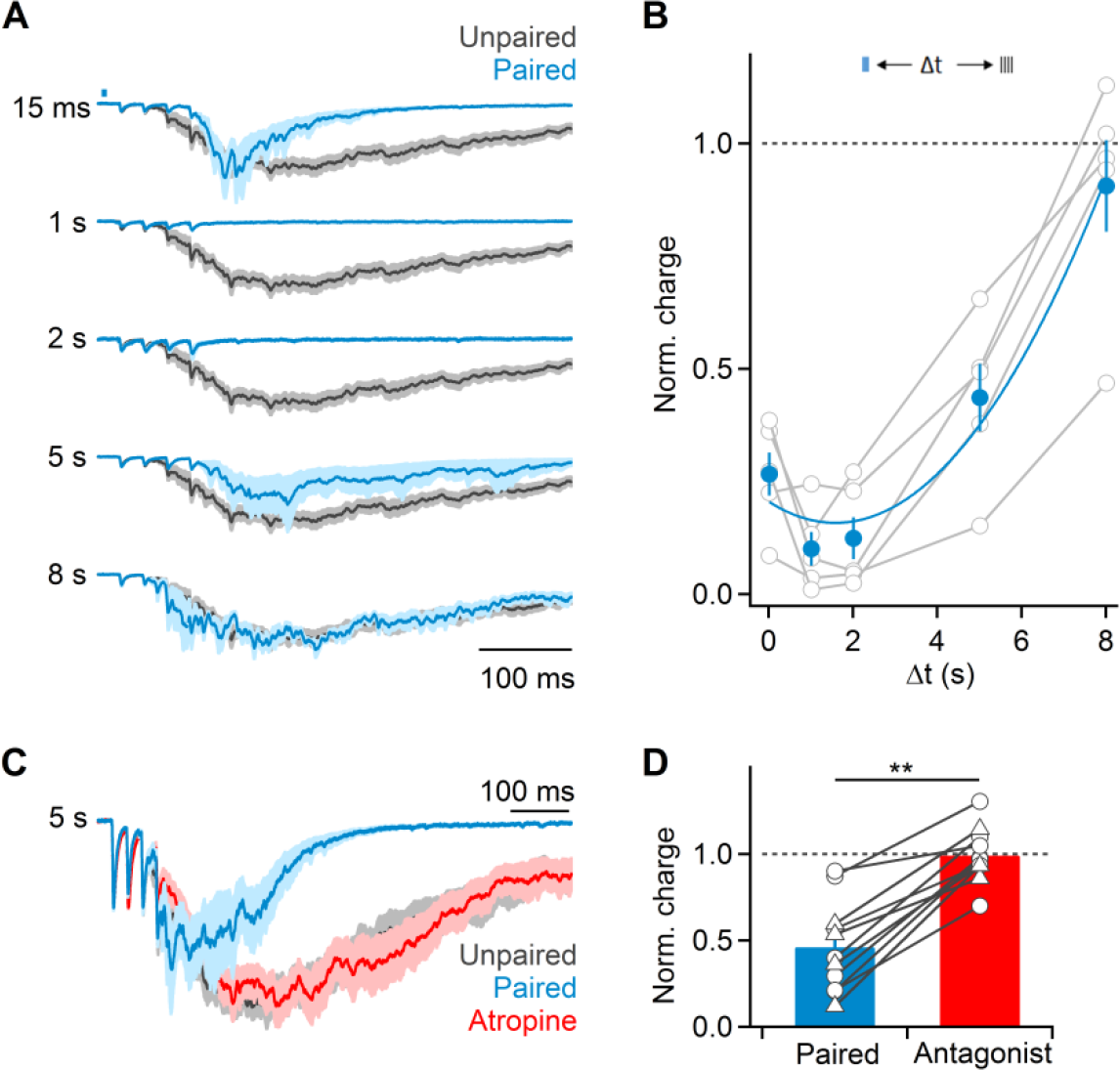
Synaptic recruitment of mAChRs mediates sustained suppression of recurrent activity. **(A)** Representative recording showing average EPSCs during unpaired (black) and paired (blue) trials, for a range of temporal delays (15 - 8000 ms) between optical and electrical stimulation. **(B)** Summary data quantifying light-evoked suppression of recurrent activity (normalized to responses in unpaired trials) as a function of temporal delay between and electrical stimulation (n = 5 cells). Summary data were fit by a third order polynomial (χ^2^=0.013). **(C)** Representative recording showing that for 5 sec delays between optical and electrical stimulation, suppression of recurrent activity (blue) was entirely reversed by bath application of atropine. **(D)** Summary data showing elimination of light-evoked suppression of recurrent activation following of bath application of either atropine or 10 µM AF-DX 116 (circles: atropine, n = 7 cells; triangles: AF-DX 116, n = 4 cells), for experiments as shown in (C). Shaded areas and error bars denote SEM.

The strong reduction of recurrent activity several seconds after the release of ACh does not appear to be compatible with a role for nAChRs. Indeed, for experiments with delays of 5 seconds between optical and electrical stimulation, bath application of atropine or the M2/M4 mAChR antagonist AF-DX 116 completely eliminated cholinergic suppression (control: 46.1 ± 8% suppression, atropine/AF: 99.2 ± 5% suppression, n = 11, p < 0.01, Wilcoxon signed rank test; Figure 3C and D). Thus, nAChRs and mAChRs mediate cholinergic suppression of recurrent activity on distinct timescales, with nAChRs mediating transient reduction and mAChRs being responsible for long-lasting reduction of cortical activity.

### Cholinergic suppression mediated by nAChRs and mAChRs is layer-specific

Next, we examined if the contributions of nAChRs and mAChRs to cholinergic suppression could be localized to distinct cortical layers. To address this question, we removed layers 1-3 by performing cuts in cortical slices just above layer 4, and carried out recordings in layer 4 (Figure 4A). Extracellular stimulation applied to the same barrel still led to recurrent activity, but with reduced magnitude (uncut slice: 105 ± 15 pC, n = 19, layer 4-6 slice: 54.9 ± 8 pC, n = 15). Furthermore, we still observed a suppression of recurrent activity (38.5 ± 5% compared to unpaired trials, n = 15, p < 0.001, Wilcoxon signed rank test; Figure 4B and C). However, in contrast to our findings in intact slices, bath application of atropine almost completely reversed cholinergic suppression (control: 35.4±7% compared to unpaired trials, atropine: 92.8 ± 4%, n = 6, p=0.01, Wilcoxon signed rank test; Figure 4B and C), suggesting that the contribution of nAChRs to controlling network activity was limited to the superficial layers. In agreement, bath application of MLA and DHβE to block nAChRs had no effect on cholinergic suppression (control: 35.1 ± 6% compared to unpaired trials, MLA and DHβE: 30.2 ± 5%, n = 6, p = 0.23, Wilcoxon signed rank test; Figure 4D and E), contrary to our observations in intact slices (Figure 2C). Furthermore, increasing the delay between optical and extracellular stimuli to 5 seconds still led to atropine-sensitive suppression of recurrent activity (control: 46.4 ± 8% compared to unpaired trials, atropine: 124.6 ± 27%, n = 5, p = 0.02, Wilcoxon signed rank test; Figure 4F and G). To further constrain the location of mAChR-mediated suppression, we performed horizontal cuts just below layer 4, and evoked recurrent activity using electrodes placed in the white matter, below the recording site in layer 5. The magnitude of recurrent activity was further reduced under these conditions (uncut slice: 105 ± 15 pC, layer 5-6 slice: 12.8 ± 3 pC, n = 6). Importantly, optical stimulation had no effect on recurrent activity (90.9 ± 8% compared to unpaired trials, n = 6, p = 0.12, Wilcoxon signed rank test; Figure 4H and I), suggesting that fast synaptic ACh release in the infragranular layers is not involved in the control of cortical activity, at least under our experimental conditions.

**Figure 4:**
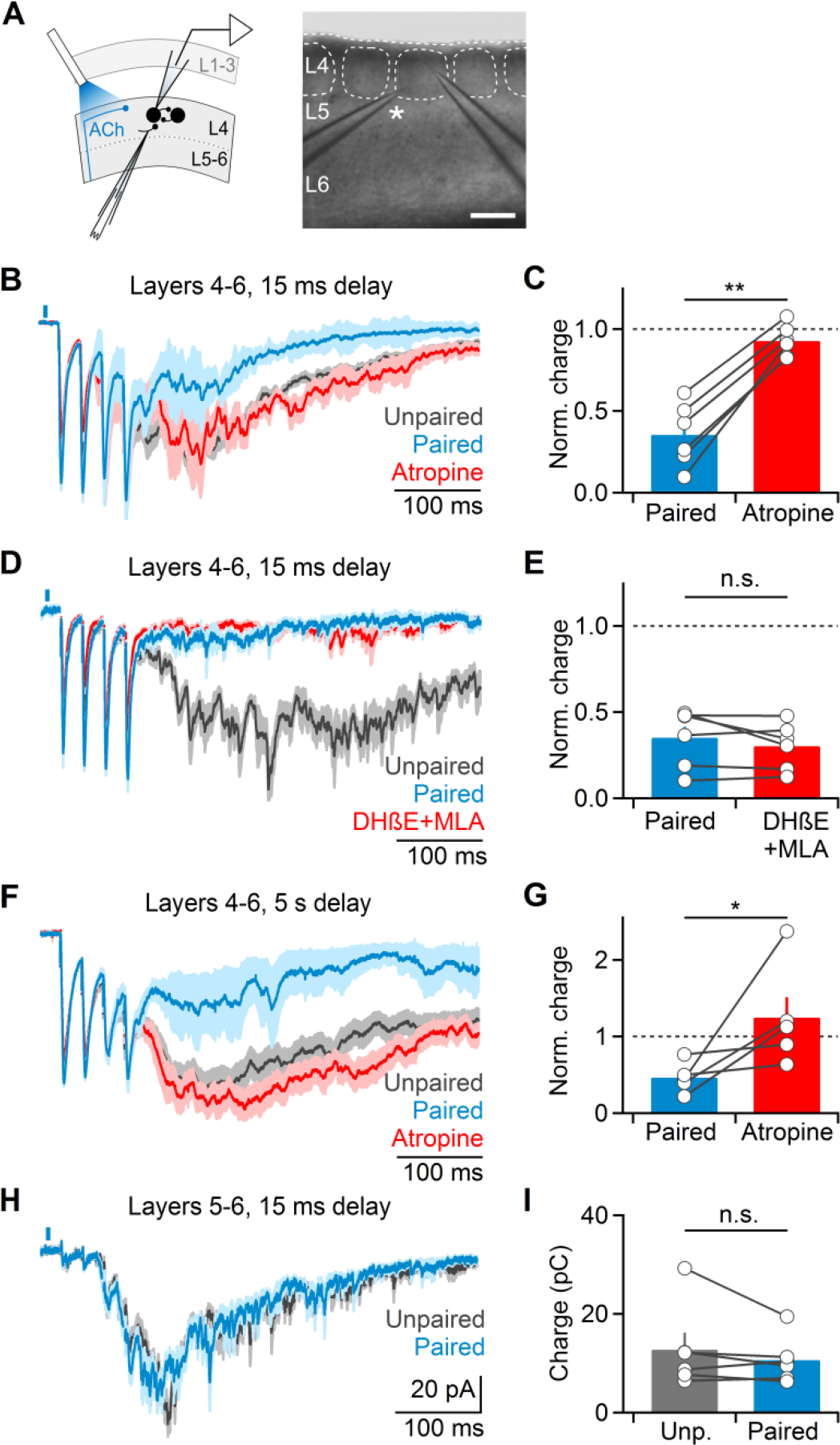
Cholinergic suppression of recurrent activity is layer-specific. **(A-G)** Recordings were carried out in slices following surgical removal of layers 1-3. **(A)** Left: Schematic indicating recording arrangement. Right: brightfield image of slice preparation. Asterisk denotes stimulating electrode. Scale bar: 150 µm. **(B)** Representative recording of layer 4 neuron showing that cholinergic suppression of recurrent activity is entirely mAChR-mediated. **(C)** Summary data (n = 6 cells) showing complete reversal of cholinergic suppression following atropine application. (**D**) Representative recording showing that nAChR antagonist application no longer reduces cholinergic suppression (n = 4 cells, DHβE alone: n = 2, DHβE+MLA: n = 2). **(E)** Summary data (n = 5 cells) for experiments as shown in (D). **(F)** Cholinergic suppression of recurrent activity is maintained for long delays (5 s) between optical and electrical stimuli and mediated my mAChRs. **(G)** Summary data (n = 5 cells) for experiments as shown in (F). **(H-I)** Surgical removal of layers 1-4 eliminates cholinergic suppression. Recordings were carried out from layer 5 neurons and activity was evoked in the white matter below the same column. **(H)** Representative recording showing EPSCs averaged across paired and unpaired trials. **(I)** Summary data (n = 6 cells) for recordings as shown in (H). All shaded areas and error bars denote SEM.

Taken together, our findings indicate that the contributions of nAChRs and mAChRs to the suppression of network activity are not uniform across cortical layers. Instead, they suggest that nAChR-dependent suppression is limited to layers 1-3, while mAChR-mediated suppression is particularly prominent in layer 4.

### Cholinergic responses are largely nicotinic in layer 2/3 and largely muscarinic in layer 4

Our results described so far predict that endogenous ACh recruits nAChRs primarily expressed in layer 2/3 GABAergic interneurons, which in turn mediate a transient form of suppression of cortical activity. In addition, they predict a prominent recruitment of mAChRs in layer 4, leading to a long-lasting depolarization of GABAergic interneurons, a long-lasting inhibition of excitatory neurons, or both. In order to test these predictions, we carried out recordings from neurons in layers 1-4 using a K^+^-based recording solution and determined the nature and frequency of optically-evoked postsynaptic responses. Neurons were classified as either regular-spiking (RS) cells considered excitatory, or as fast-spiking (FS) or non-fast-spiking (non-FS) cells considered inhibitory, based on their intrinsic firing properties (Beierlein et al., 2003) (Figure 5A). In agreement with previous findings (Arroyo et al., 2012), neurons in layer 1 showed nAChR-mediated EPSCs (nEPSCs, 11/12 neurons) that were fully blocked by a combination of MLA and DHβE (data not shown). In layer 2/3, a large percentage of inhibitory interneurons displayed nEPSCs that were blocked by DHβE (FS: 39%, non-FS: 77%; Figure 5B and C), while a minority of neurons displayed long-lasting mAChR-dependent currents (FS: 23%, non-FS: 5%; Figure 5B and C). To confirm the existence of functional mAChRs in non-FS neurons as shown previously (Chen et al., 2015), we used a picospritzer to apply brief puffs of muscarine. For all neurons examined (n = 9) which showed an optically-evoked nEPSP only, muscarine application led to a robust depolarization which was blocked by atropine (Figure 5-figure supplement 1). These data indicate that while mAChRs are prominently expressed in layer 2/3 non-FS neurons, they are not recruited by synaptically released Ach under our conditions.

**Figure 5:**
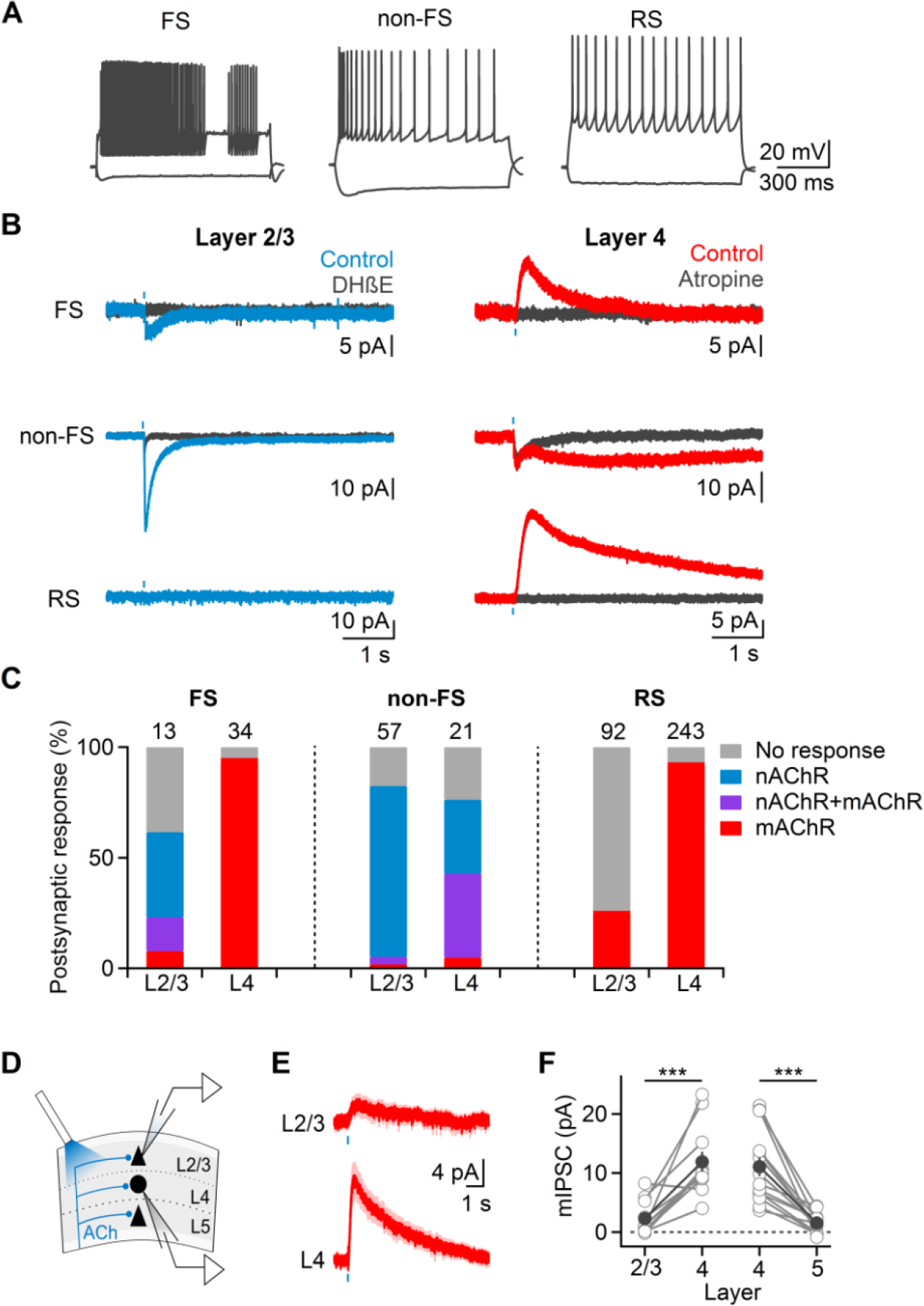
Cholinergic postsynaptic responses are layer-specific. **(A)** Cells in layer 2/3 and layer 4 were classified as either inhibitory FS, or non-FS cells or excitatory RS cells based on their intrinsic firing characteristics. **(B)** Example voltage clamp recordings carried out in the presence of NBQX (10 µM), DAPV (25 µM), picrotoxin (50 µM) and CGP 55845 (5 µM) showing typical light-evoked responses. Most cells in layer 2/3 (left column) showed either no response or fast EPSCs (blue traces) blocked by DHβE (black traces). In layer 4 (right column) the majority of neurons displayed slow postsynaptic responses (red traces) that were blocked by atropine (black traces). **(C)** Summary data showing likelihood of nAChRs and mAChR-mediated responses for each cell type. Numbers above bars indicate total number of cells recorded. **(D-F)** mIPSCs in RS cells are strongest in layer 4. **(D)** Schematic of recording setup. Simultaneous recordings of pairs of layer 4 & 2/3, or layer 4 & 5 RS cells were obtained. Postsynaptic responses in each cell were evoked with optical stimulation centered over respective neuron **(E)** Average IPSCs recorded across pairs of layer 2/3 and layer 4 neurons (n = 11 pairs). **(F)** Summary data of dual recordings (11 layer 2/3 & 4 pairs, 14 layer 4 & 5 pairs). All shaded areas and error bars denote SEM.

In contrast to interneurons, most RS cells in layer 2/3 did not show optically-evoked postsynaptic responses (75%; Figure 5B and C), with the remaining neurons displaying small-amplitude mAChR-dependent IPSCs (mIPSCs, 25%). Taken together, these findings indicate that the synaptic release of ACh in superficial layers controls cortical activity primarily via the recruitment of nAChRs in distinct types of interneurons.

Recordings in layer 4 yielded dramatically different results. The vast majority of FS cells displayed atropine-sensitive mIPSCs (94%; Figure 5B and C), and never showed nAChR dependent responses. Non-FS interneurons responding to ACh release displayed either isolated nEPSCs (33%), or biphasic responses consisting of nEPSCs and mAChR EPSCs (mEPSCs, 38%; Figure 5B and C). Surprisingly, the large majority of RS cells showed mIPSCs (92%) that were blocked by atropine or AF-DX 116. Postsynaptic mAChR-dependent responses displayed large cell type-specific differences in their kinetics (Figure 5-figure supplement 2). While mIPSCs in FS had relatively fast kinetics (rise time: 165.9 ± 10 ms, decay time constant: 844.2 ± 78 ms, n = 20), mIPSCs in RS cells were considerably slower (rise time: 328.3 ± 23 ms, decay time constant: 3281.7 ± 157 ms, n = 21, p<10^−5^, ANOVA). Finally, mEPSCs in non-FS cells displayed extremely slow kinetics (rise time: 1248.3 ± 125 ms, decay time constant: 23.7 ± 5.4 s, n = 7, p < 10^−5^, ANOVA). Thus, the recruitment of mAChRs by synaptically released ACh can control postsynaptic activity on dramatically different time scales, depending on the cell type.

Our data suggest that excitatory neurons in layer 4 are much more likely to receive cholinergic inputs compared to excitatory neurons in layers 2/3. To better quantify the functional impact of cholinergic innervation to excitatory neurons located in distinct layers while accounting for variable ChR2 expression levels in different animals, we performed dual recordings from RS neurons in layer 4 and layer 2/3 or layer 4 and layer 5 in the same cortical column (Figure 5D). For all pairs examined, mIPSC amplitudes in layer 4 were significantly larger compared to responses in either layer 2/3 or layer 5 (layer 2/3: 2.3 ± 1 pA vs. layer 4: 11.9 ± 2 pA, n = 11 pairs, p < 0.001; layer 5: 1.5 ± 0 pA vs. layer 4: 11 ± 2 pA, n = 14 pairs, p < 0.0001, two-tailed paired t-test; Figure 5E,F), further confirming that the mAChR-dependent cholinergic signaling is most robust in layer 4.

Next, we probed the mechanisms mediating mIPSCs in layer 4 RS neurons. Synaptic currents had onset latencies of 30.6 ± 1 ms (n = 19 cells), reversed at ~-96 mV, displayed strong inward rectification and could be blocked by bath application of barium (control: 9.9 ± 3 pA, Ba^2+^: 1.6 ± 1 pA, n = 4 cells; Figure 5-figure supplement 3), indicating the mIPSCs were mediated by G protein-coupled inwardly-rectifying potassium (GIRK) conductances. By contrast, bath application of the small conductance calcium-activated potassium (SK) channel antagonist apamin had little effect on mIPSC amplitudes (control: 12 ± 2 pA, apamin: 10.9 ± 1 pA, n = 6 cells; Figure 5-figure supplement 3) and recordings using an internal solution containing 5 mM BAPTA did not attenuate mIPSCs (n = 4 cells, data not shown), suggesting that SK channel activation is not involved in mediating mIPSCs in layer 4 neurons.

Taken together, our data show that the synaptic release of ACh in layer 4 leads to the recruitment of mAChRs in all major cell types including the large majority of RS cells, suggesting that direct cholinergic inhibition of excitatory neurons plays a central role in the prolonged suppression of recurrent activity.

### Synaptic ACh reduces neuronal firing in layer 4 RS cells via hyperpolarizing inhibition

Next, we quantified the functional impact of the fast activation of mAChRs in layer 4 RS cells. For these experiments, we paired action potential activity evoked by depolarizing current steps with optical activation of cholinergic afferents. For all neurons tested (n = 11 cells), postsynaptic firing frequencies following optical stimuli were rapidly (<100 ms) and persistently reduced compared to control trials (Figure 6A-C). To determine the functional impact of cholinergic synaptic signaling on the integration of synaptic inputs, we paired optical stimulation with action potential activity evoked by extracellular stimulation of glutamatergic afferents (4 stimuli at 40 Hz), under conditions where recurrent activity was blocked pharmacologically with the NMDAR antagonist APV (25 µM) (Beierlein et al., 2002). Light-evoked mAChR IPSPs (mIPSPs) delayed or blocked action potential activity compared to unpaired trials (unpaired: 1.4 ± 0.1 spikes per trial, paired: 0.87 ± 0.1 spikes per trial, n = 10 cells, p<0.01, Wilcoxon signed rank test; Figure 6-figure supplement 1). Together, these data demonstrate that the synaptic activation of mAChRs leads to a rapid and long-lasting reduction of RS cell activity in layer 4, consistent with a critical role in reducing recurrent activity.

**Figure 6:**
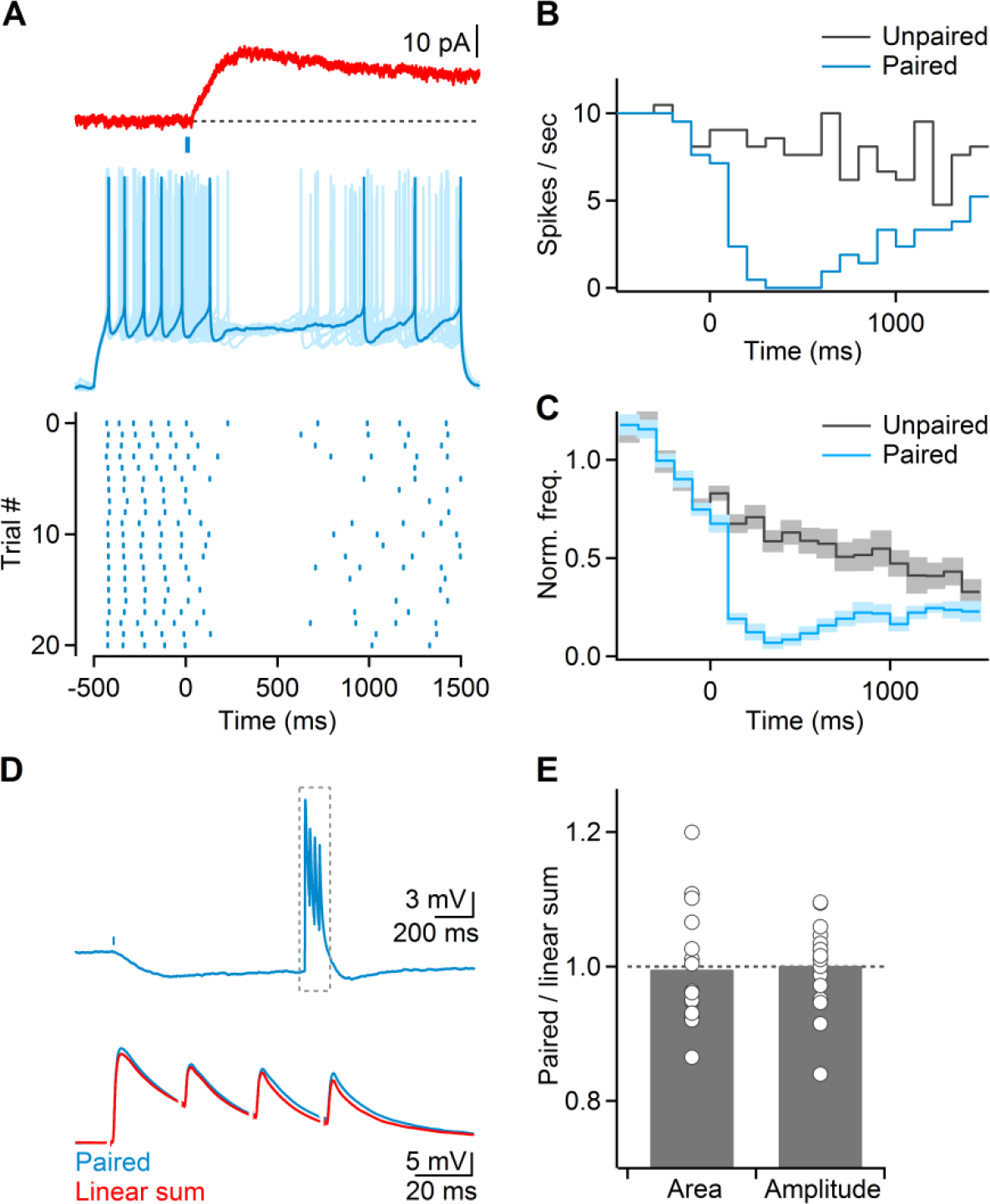
Cholinergic afferents control RS cells in layer 4 via hyperpolarizing inhibition. **(A)** *Top:* mIPSC in layer 4 RS cell. *Middle:* recordings in multiple trials showing current-step evoked neuronal activity paired with light stimulation (indicated by blue bar). B*ottom:* Raster plot indicating action potential timing over multiple trials, for the same neuron. **(B)** For the same cell as in (A), peristimulus time histogram (PSTH) for paired (blue) and unpaired (gray) current steps. **(C)** Average PSTH (n = 11 RS cells), normalized to average of first five 100 ms time bins for each cell. **(D)** mAChR-mediated IPSPs (mIPSP) do not cause shunting of glutamatergic EPSPs. *Top*: optically evoked mIPSP was paired with a train (40 Hz) of electrically evoked glutamatergic EPSPs (delay: 1 sec). Recordings were carried out in the presence of D-APV (25 µM) to prevent recurrent activity. *Bottom*, close-up of recording for time window indicated in top trace showing that paired response (blue trace) is identical to linear sum of mIPSP and EPSPs evoked separately (red trace). **(E)** Summary data quantifying both area under the paired EPSPs and amplitude of the first paired EPSP, normalized to their respective unpaired controls (n = 16 cells).

By what mechanism do mIPSPs influence synaptic integration and neuronal firing in excitatory neurons? Activation of mAChRs and the opening of GIRK conductances leads to a hyperpolarization of membrane potential and in addition, to an increase in membrane conductance generating a potential “shunt” (Eggermann and Feldmeyer, 2009). Shunting inhibition is thought to be a prominent mechanism underlying the spatiotemporal summation of excitatory and inhibitory synaptic inputs in neocortex and other brain areas (Koch, 1999). To determine if cholinergic afferents generate shunting inhibition in layer 4 RS neurons, we simultaneously evoked mIPSPs and subthreshold glutamatergic EPSPs. Surprisingly, we found that both glutamatergic EPSP amplitude and area of the paired postsynaptic response were on average nearly identical to the linear sum of the EPSP and the mIPSP evoked separately (EPSP amplitude: 100.1 ± 2% compared to linear sum, n = 16, p = 0.16, EPSP area: 99.6 ± 2%, n = 16, p = 0.34, Wilcoxon signed rank test; Figure 6D and E), suggesting that shunting inhibition was not prominent. Instead, our data indicate that cholinergic inputs to layer 4 excitatory cells reduce neuronal firing primarily via hyperpolarizing inhibition.

## Discussion

Previous work performed in vivo and in vitro has provided extensive evidence that the fast activation of GABAergic interneurons by nAChRs can mediate cholinergic control of cortical activity. However, the conditions leading to the recruitment of mAChRs are far less understood. Here we have demonstrated that the release of ACh evoked by individual pulses can reliably activate postsynaptic mAChRs in the large majority of excitatory neurons of layer 4, ultimately leading to a strong suppression of cortical activity lasting several seconds. Our results highlight a critical role of mAChRs in the rapid and flexible modulation of cortical brain state.

### Mechanisms underlying cholinergic synaptic signaling

We found that activation of cholinergic afferents led to mAChR-dependent IPSCs in excitatory neurons of layer 4. IPSCs were blocked by the M2/M4 specific antagonist AF-DX 116, showed strong inward rectification and were blocked by barium, consistent with the activation of M2/M4 mAChRs leading to GIRK opening by a membrane delimited pathway, as was observed previously for thalamic cholinergic synapses (Sun et al., 2013). This extends previous findings in layer 4 of rat barrel cortex showing GIRK activation following exogenous agonist application (Eggermann and Feldmeyer, 2009). At the same time, we did not find evidence for an involvement of SK channels in generating mAChR-dependent inhibition, in contrast to studies employing exogenous agonist application in other cortical areas and layers (Gulledge and Stuart, 2005; Gulledge et al., 2007).

Surprisingly, mAChR-dependent postsynaptic responses in layer 4 RS cells had no effect on the amplitude of glutamatergic EPSPs, i.e. they did not generate shunting inhibition, contrary to results obtained using bath application of agonists (Eggermann and Feldmeyer, 2009). Besides being strongly distance-dependent, shunting inhibition is most effective for inhibitory synapses located “on-path“, i.e., between excitatory synapses and the spike initiation zone in the axon (Vu and Krasne, 1992). Since our extracellular stimuli likely recruited both intracortical and thalamocortical glutamatergic synapses targeting the entire dendritic arbor, cholinergic synapses are predicted to be located in distal dendritic regions. Alternatively, cholinergic transmission might lead to the activation of multiple postsynaptic signaling cascades, resulting ultimately in little overall change of membrane conductance. How cholinergic synaptic signals dynamically regulate postsynaptic integration in dendritic trees will require further investigation.

What is the signaling mode responsible for the synaptic recruitment of mAChRs in layer 4? It remains unclear how widespread ultrastructurally defined cholinergic synapses are in neocortex, with studies reporting either classic synapses (Turrini et al., 2001), small varicosities (Umbriaco et al., 1994) or a complete absence of distinct release sites. Furthermore, specific mAChR subtypes do not appear to be preferentially expressed near suspected sites of ACh release (Yamasaki et al., 2010), suggesting that mAChRs only respond to slow increases in cortical ACh levels following sustained increases in BF afferent activity that can lead to transmitter spillover (Descarries et al., 1997). However, such a scenario is difficult to reconcile with the relatively low firing rates of BF cholinergic neurons, even during awake states (Lee et al., 2005; Hassani et al., 2009; Hangya et al., 2015). Our demonstration of short-latency (~30 ms) mAChR-mediated responses in the large majority of layer 4 neurons suggests that mAChRs can participate in point-to-point transmission, via either conventional synapses or unique forms of volume transmission, as has been suggested for certain types of nAChR-mediated responses (Bennett et al., 2012). However, our findings also show that extrasynaptic mAChRs in layer 2/3 non-FS neurons and presynaptic mAChRs are not readily recruited by brief activation of cholinergic afferents, suggesting the existence of slower and more diffuse forms of signaling in need of further investigation.

### The role of interneurons in cholinergic control of cortical activity

Many questions remain regarding the role of interneurons in mediating cholinergic control of cortical processing. Elegant in vivo studies have shown that cholinergic afferents can rapidly engage layer 1 interneurons (Letzkus et al., 2011) as well as vasoactive intestinal peptide (VIP)-expressing interneurons (Fu et al., 2014) by activating postsynaptic nAChRs. As both types of neurons primarily target other interneurons (Lee et al., 2013; Pi et al., 2013) which in turn project onto pyramidal neurons, cholinergic inputs in superficial cortical layers can increase activity in cortical networks by disinhibition (Letzkus et al., 2015). Here we find that nAChR-mediated cholinergic signaling in fact moderately *suppresses* cortical activity. This indicates that at least under our experimental conditions, cholinergic inputs in superficial cortical layers increase activity in interneurons that preferentially target excitatory cells, including FS and somatostatin-expressing (SOM) non-FS cells, ultimately generating feed-forward inhibition. The balance between BF-evoked inhibition and disinhibition is likely to be highly dependent on the activity levels of distinct types of interneurons associated with different cortical activity patterns (Moore et al., 2010).

Contrary to our findings in layer 2/3, a high percentage of layer 4 non-FS neurons displayed mAChR-mediated depolarizations, consistent with recent in vivo results (Muñoz et al., 2017). Therefore, it is possible that mAChR-mediated excitation of layer 4 non-FS interneurons is at least partly responsible for cholinergic suppression. However, because mAChR-mediated EPSCs in these cells show extremely slow rise times, they are unlikely to contribute to cholinergic suppression at short latencies. Furthermore, the majority of non-FS cells we recorded from in layer 4 are likely to be SOM neurons (Rudy et al., 2011) and activation of these cells is predicted to generate *disinhibition* of layer 4 cortical activity, via their preferential inhibition of FS cells (Xu et al., 2013). Taken together, the mAChR-dependent suppression of network activity we observed is likely to be largely mediated by a direct monosynaptic inhibition of excitatory neurons.

### Cholinergic control of cortical state

Information processing in cortical circuits is strongly modulated by the internal cortical state, as defined by the degree of low-frequency synchronous activity in the local network (Harris and Thiele, 2011). Transitions between cortical states can have dramatically different temporal dynamics, ranging from very slow and sustained as for sleep-wake transitions, to very rapid and highly transient, as observed for sub-states within the awake state (McGinley et al., 2015b). Fast state transitions from quiet wakefulness to a medium arousal state or to locomotor behavior are characterized by a suppression of low frequency rhythmic activity, thereby enabling increased sensory responses and improved behavioral performance. The mechanisms underlying fast cortical state transitions remain poorly understood but likely involve a number of highly coordinated processes, including several distinct neuromodulatory systems (Aston-Jones and Cohen, 2005; McGinley et al., 2015b) and changes in thalamocortical and corticothalamic activity patterns (Zagha and McCormick, 2014). BF cholinergic afferent activity is likely critical for mediating moment-to-moment changes in brain state (Eggermann et al., 2014; Reimer et al., 2016). For example, BF afferents to neocortex increase their activity prior to whisking and BF activity is causally linked to a suppression of low frequency oscillations for several seconds typical for quiet wakefulness (Eggermann et al., 2014).

Our findings confirm previous studies showing a reduction of spontaneous slow oscillatory cortical activity following exogenous ACh application (Favero et al., 2012; Wester and Contreras, 2013; Castro-Alamancos and Gulati, 2014). In addition, they offer a physiologically plausible mechanism for the precise spatiotemporal control of cortical activity by synaptically released ACh. In our hands, even brief ACh transients were sufficient to cause a prolonged suppression of low-frequency cortical activity, via the rapid activation of mAChRs. Furthermore, our findings suggest that the long-lasting hyperpolarization of layer 4 excitatory neurons is a critical substrate of cholinergic action. This implies that the initial processing of thalamic inputs is under direct cholinergic control, with cholinergic afferent activity tracking increases and decreases in thalamic afferent activity associated with different behavioral states. Such ongoing adjustments in layer 4 gain might underlie low-noise cortical computations during periods of heightened arousal. In addition, long-lasting gain control in layer 4 might allow for more rapid nAChR-mediated computations involving both inhibition and disinhibition in local circuits in superficial layers (Letzkus et al., 2015).

## Materials and Methods

### Slice preparation

Recordings were obtained in acute brain slices prepared from both male and female bacterial artificial chromosome (BAC)-transgenic mice expressing ChR2 under the control of the choline acetyltransferase (ChAT) promoter (ChAT–ChR2–EYFP; (Zhao et al., 2011)). Some experiments were performed in slices derived from C57BL/6 wild-type mice. All animals used in this study were treated following procedures in accordance with National Institutes of Health guidelines and approved by the University of Texas Health Science Center at Houston (UTHealth) animal welfare committee. Animals aged P12-16 were anesthetized using isoflurane and then decapitated. The brains were rapidly removed and placed in ice cold cutting solution saturated with 95% O_2_–5% CO_2_, that consisted of the following (in mM): 212 sucrose, 2.5 KCl, 1.25 NaH_2_PO_4_, 10 MgSO_4_, 0.5 CaCl_2_, 26 NaHCO_3_, and 11 glucose. Thalamocortical slices (Agmon and Connors, 1991) (400 µm) were cut using a VT1200 S vibratome (Leica) and immediately transferred to artificial cerebrospinal fluid (ACSF, saturated with 95% O_2_–5% CO_2_), maintained at 35°C and consisting of the following (in mM): 126 NaCl, 2.5 KCl, 1.25 NaH_2_PO_4_, 2 MgCl_2_, 2 CaCl_2_, 26 NaHCO_3_ and 10 glucose. Slices were incubated at 35°C for 20 minutes and then stored at room temperature until used for experiments.

### Recording

Electrophysiological recordings were performed in a recording chamber perfused with ACSF saturated with 95% O_2_–5% CO_2_ and warmed to 31-34°C using a Warner Instruments TC-324B in-line heater. Cells were visualized via infrared differential interference contrast (IR-DIC) under an Olympus BX51WI microscope equipped with a Dage-MTI IR-1000 camera. Extracellular electrical stimuli (1-20 µA) were delivered via a glass electrode filled with ACSF, using an A-M Systems Model 2100 pulse stimulator. Cholinergic afferents were activated via 5 ms pulses of blue light delivered through a 60x water-immersion objective using a Prizmatix LED. Whole-cell recordings were obtained using glass pipettes with a resistance of 3-5 MΩ. For voltage clamp recordings of recurrent activity, recording pipettes were filled with a cesium-based internal solution consisting of (in mM): 120 CsMeSO_3_, 1 MgCl_2_, 1 CaCl_2_, 10 CsCl, 10 HEPES, 3 QX-314, 11 EGTA, 2 Mg-ATP, and 0.3 Na-GTP (adjusted to 295 mOsm and pH 7.3). For current clamp recordings, and voltage clamp recordings of cholinergic postsynaptic responses, we used a potassium-based internal solution consisting of (in mM): 133 K-Gluconate, 1 KCl, 2 MgCl_2_, 0.16 CaCl_2_, 10 HEPES, 0.5 EGTA, 2 Mg-ATP, and 0.4 Na-GTP (adjusted to 290 mOsm and pH 7.3). Where indicated, 5 mM BAPTA was included to block increases in intracellular calcium concentration. Muscarine was applied using a Picospritzer (Parker Automation) connected to glass electrodes that were filled with 1 mM muscarine.

NBQX, DHβE, AF-DX 116, picrotoxin, CGP 55845, D-APV, and MLA were obtained from R&D Systems. All other chemicals were obtained from Sigma-Aldrich.

### Data acquisition and analysis

Recordings were acquired using a Multiclamp 700B amplifier (Molecular Devices), filtered at 3– 10 kHz, digitized at 20 kHz with a 16-bit analog-to-digital converter (Digidata 1440A; Molecular Devices). All data were saved using Clampex 10.3 software (Molecular Devices) and analyzed using custom macros written in IGOR Pro (Wavemetrics). Statistical analyses were performed in Prism 5 software (Graphpad).

Evoked recurrent activity was quantified as total charge transferred to the recorded cell. For voltage clamp PSC recordings, we calculated the area under the PSC trace in a time window starting 90±3 ms after the first electrical pulse, and ending when evoked activity returned to baseline. For a given cell, the same time window was used for paired and unpaired trials. For pharmacological experiments, data from paired trials in a given drug condition were normalized to their respective unpaired trials in the same condition.

## Acknowledgements

This work was supported by a National Institute of Neurological Disorders and Stroke grant R01 NS077989 to M.B. and a Zilkha Family Discovery Fellowship in Neuroengineering to R.D.

---

**Figure 1-figure supplement 1:**
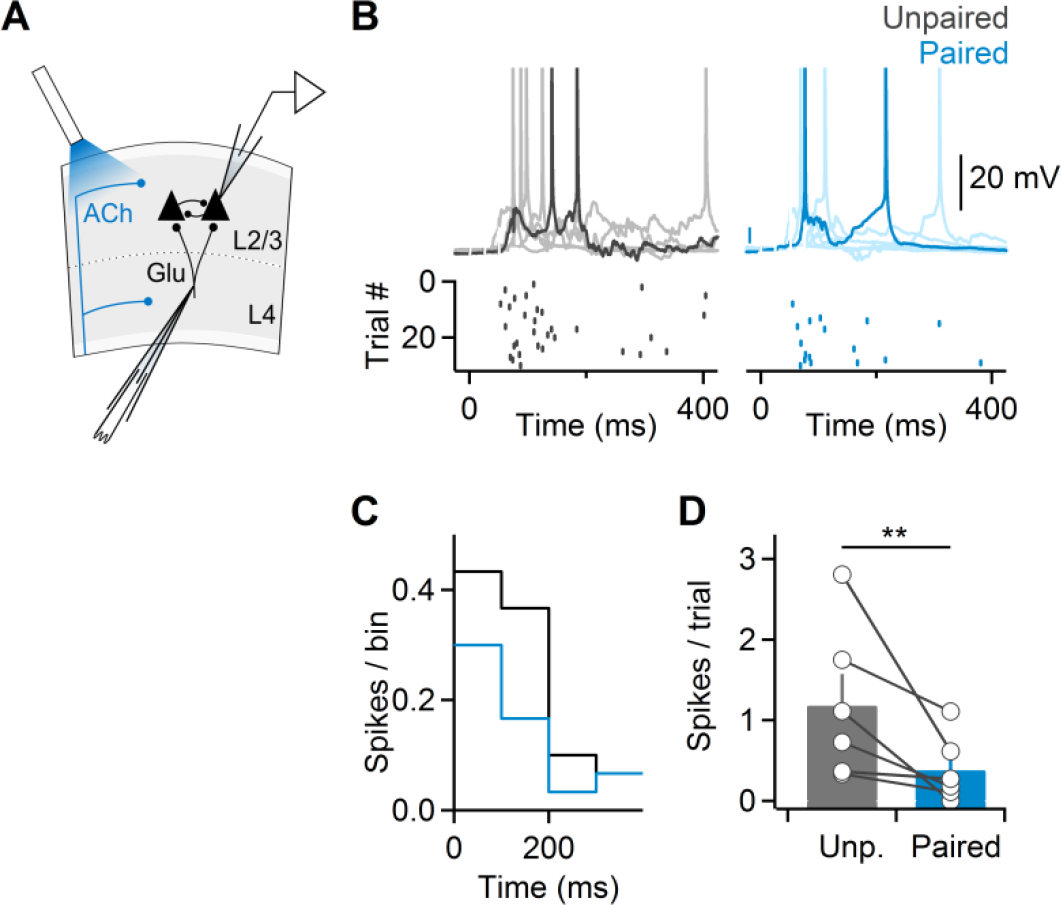
ACh release reduces action potential firing during recurrent activity. **(A)** Schematic of experimental setup. Cells were held in current clamp with minimal current injection to keep the cell at ~-60 mV. For unpaired trials, recurrent activity evoked via electrical stimulation in layer 4 was monitored in layer 2/3 neurons. For paired trials, cholinergic afferents were activated with individual light pulses (5 ms duration), 15 ms prior to electrical stimulation in layer 4. **(B)** Representative experiment showing consecutive trials of recurrent activity, under paired (black) or unpaired (blue) conditions. Raster plots indicate the timing of action potentials in individual trials. **(C)** Peristimulus time histogram for the same cell showing total spikes per 100 ms bin across trials. (**D**) Summary data (n = 6 layer 2/3 neurons) showing ACh-mediated decrease in neuronal activity.

**Figure 1-figure supplement 2:**
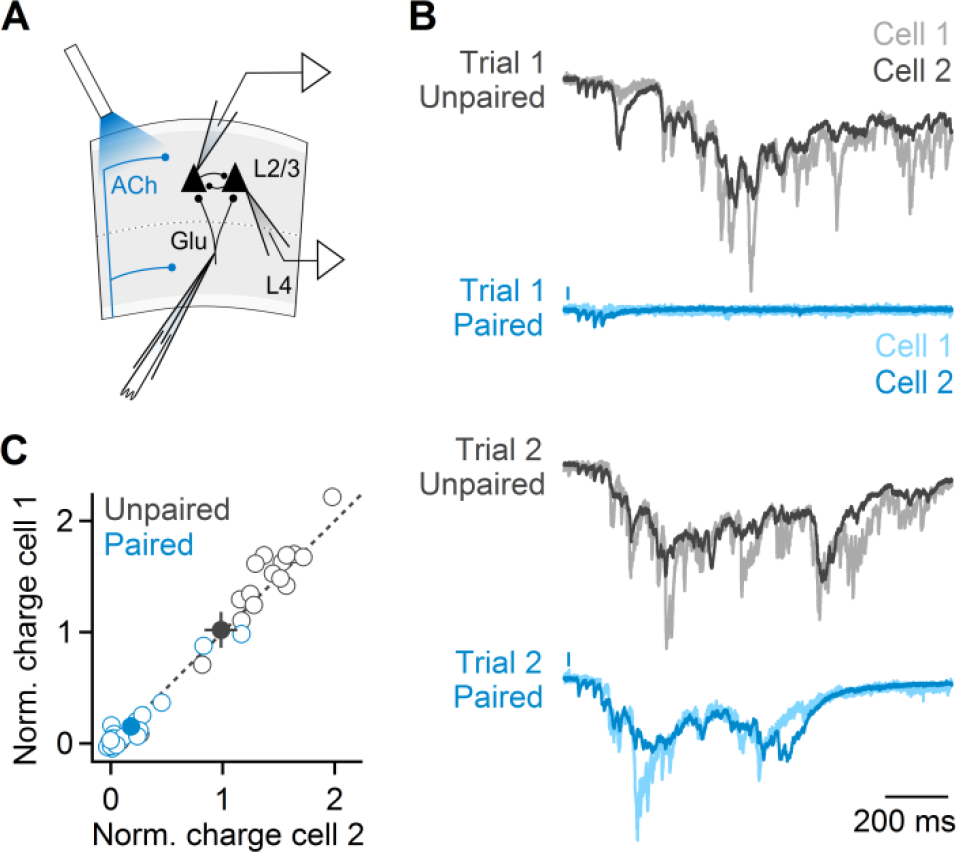
Magnitude and suppression of recurrent activity are tightly correlated within layer 2/3 local networks. **(A)** Recurrent activity evoked in layer 4 was recorded in voltage clamp from two neighboring layer 2/3 neurons (held at −70 mV to isolate EPSCs), with (paired) or without (unpaired) prior activation of cholinergic afferents. **(B)** Representative experiment showing overlaid responses from both neurons for two paired (blue) and two unpaired (black) trials recorded consecutively. **(C)** Magnitude of recurrent activity (measured as EPSC charge transfer) in individual trials (n = 22 trials) for the two cells shown in (B). Responses for each cell are normalized to the respective average recurrent activity across all unpaired trials. Filled circles denote average responses. Error bars denote SEM.

**Figure 5-figure supplement 1:**
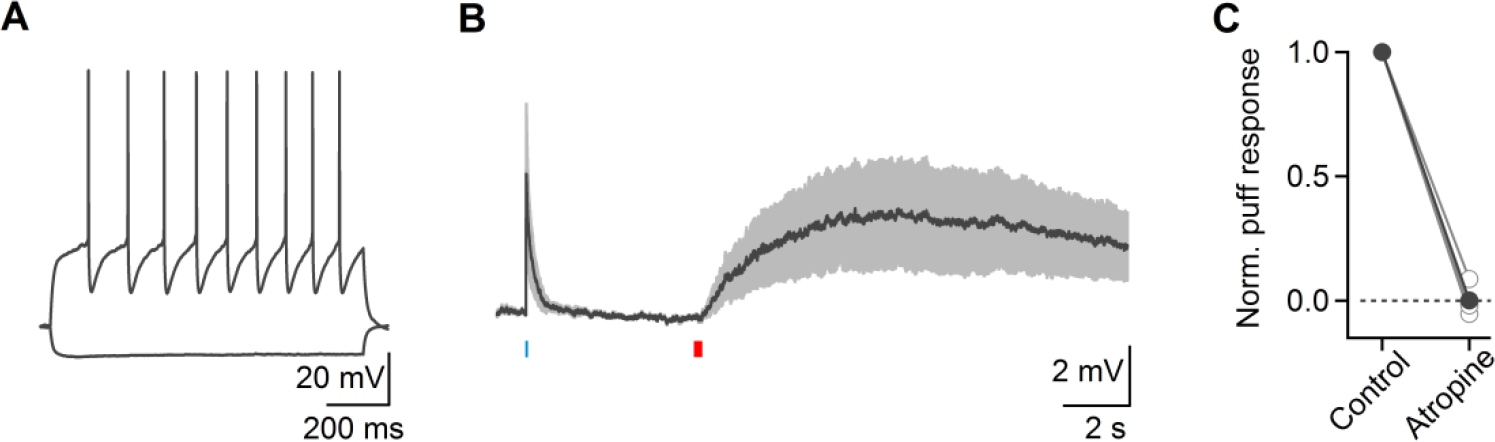
Synaptic release of ACh primarily recruits nAChRs in layer 2/3 nonFS neurons. **(A)** Responses to hyperpolarizing and depolarizing current steps in an example layer 2/3 non-FS neuron. **(B)** Average response in current clamp from layer 2/3 non-FS cells (n = 4) to 5 ms optical activation (blue bar) followed by a 200 ms puff of muscarine chloride (1 mM, red bar). **(C)** Summary data (n = 4 cells), showing that responses evoked by muscarine were completely blocked by atropine. Shaded areas and error bars denote SEM.

**Figure 5-figure supplement 2:**
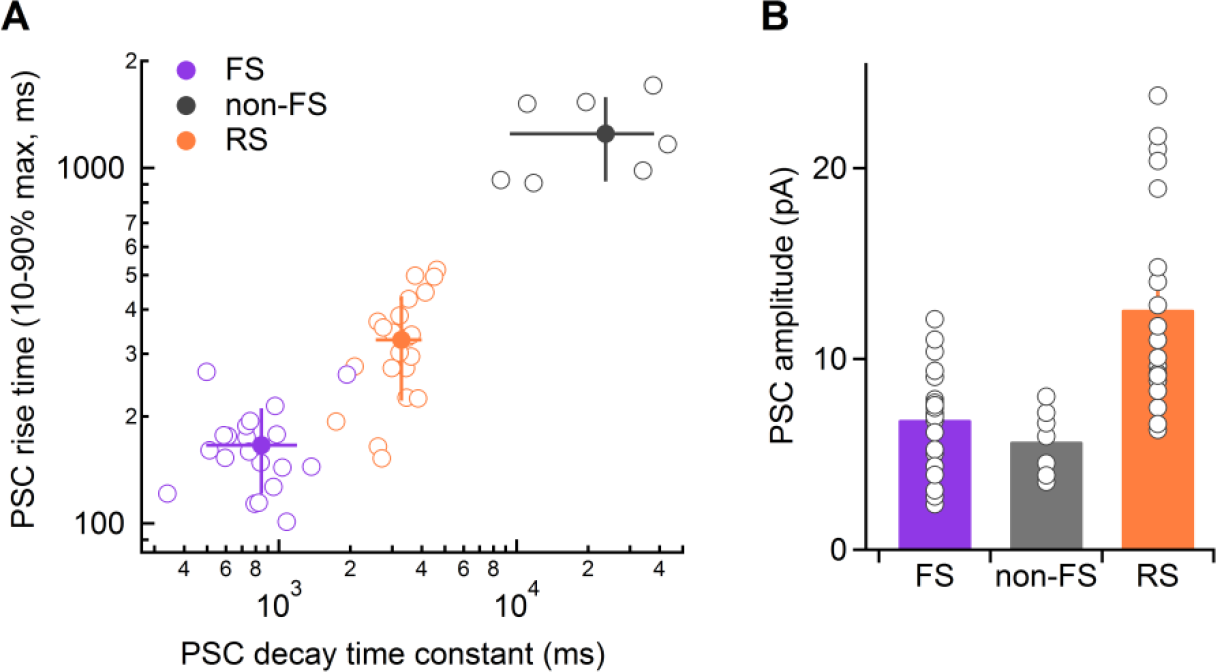
Postsynaptic mAChR-mediated responses in layer 4 have cell-type specific kinetics. **(A)** Rise times of mIPSCs plotted against their decay time constants from example layer 4 RS (n = 21) and FS (n = 20) cells, along with the same values for mEPSCs in layer 4 non-FS cells (n = 7). Note logarithmic scale on both axes. Error bars denote SD. **(B)** Summary of mAChR-mediated postsynaptic responses for the same cells as in (A). Error bars denote SEM.

**Figure 5-figure supplement 3:**
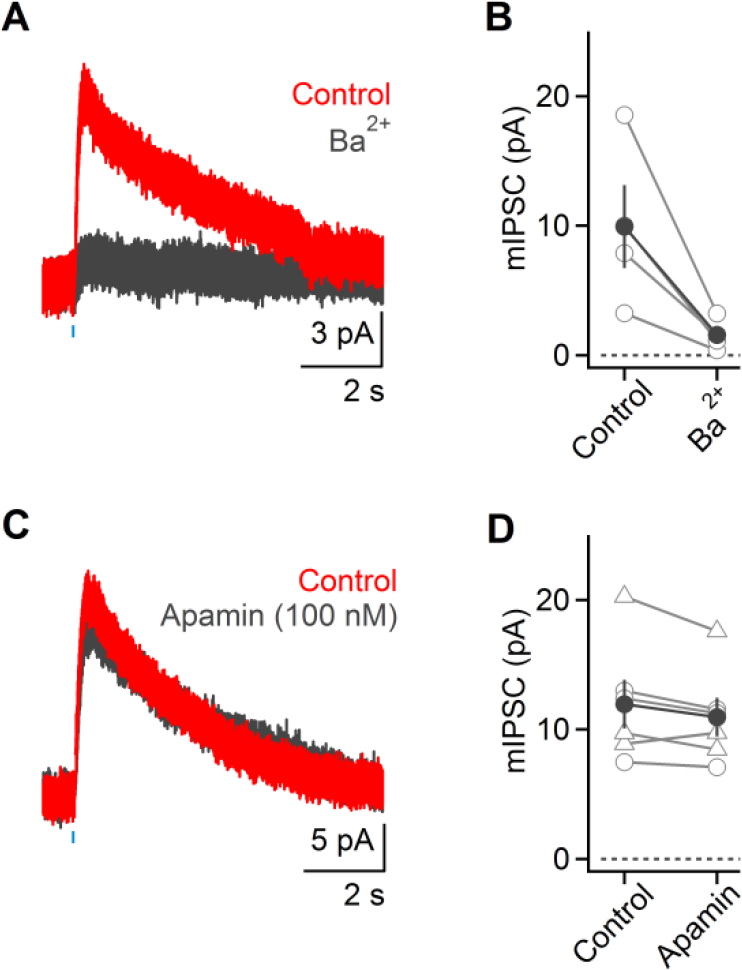
mIPSCs in layer 4 RS cell are mediated by GIRK conductances. Recordings were obtained in the presence of antagonists for GABAergic and glutamatergic synaptic transmission. **(A)** Light-evoked mIPSC in a layer 4 RS cell was blocked following Ba^2+^ (200 µM) application. **(B)** Summary data quantifying mIPSC reduction following Ba^2+^ application (n = 4 cells). **(C)** Application of the SK channel antagonist apamin (10 – 100 nM) has no effect on mIPSCs, as shown for this layer 4 RS neuron. **(D)** Summary data quantifying mIPSC responses prior to and following apamin wash-in (circles: 10 nM, n = 3 cells; triangles: 100 nM, n = 3 cells). All error bars denote SEM.

**Figure 6-figure supplement 1:**
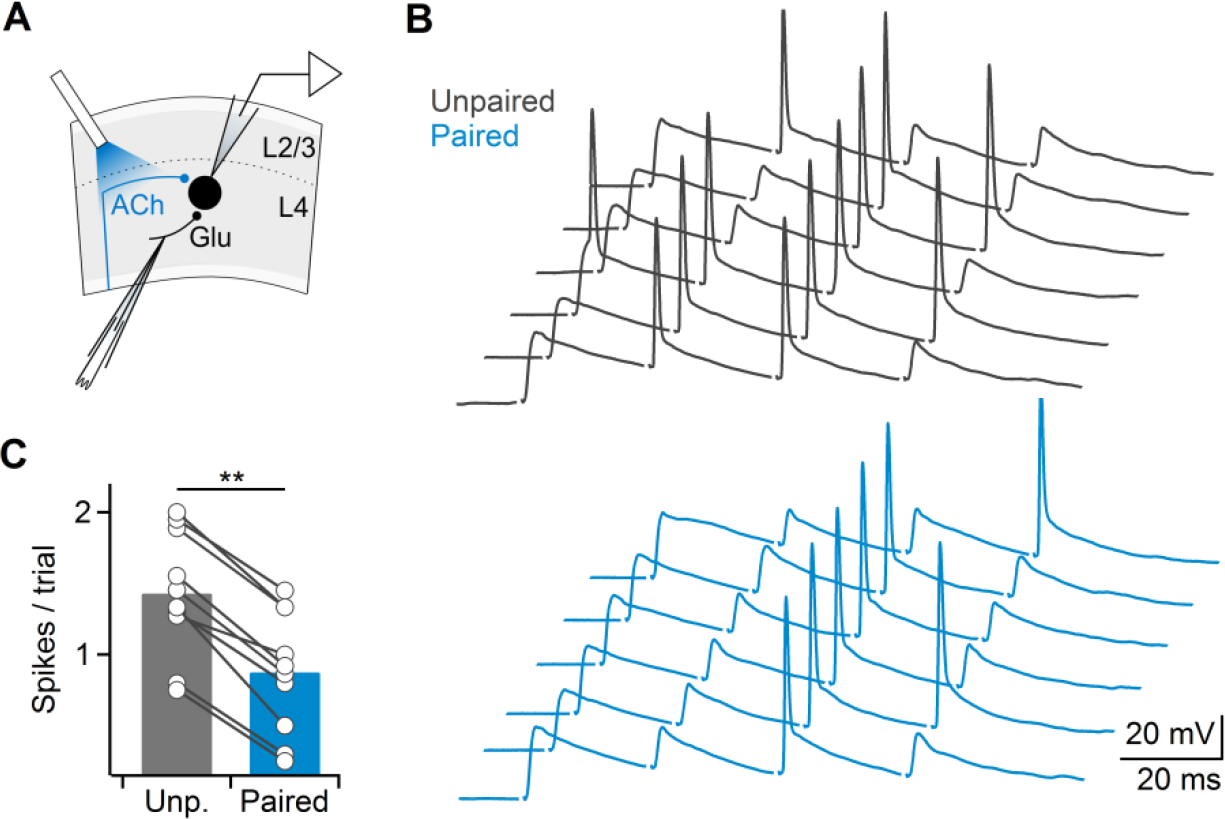
Cholinergic activation suppresses synaptically-evoked spiking in layer 4 RS cells. **(A)** Schematic of experimental setup. Glutamatergic EPSPs in layer 4 neuron were paired with single optical stimulus (5 ms), applied 1 s prior to electrical stimulation. **(B)** Glutamatergic-evoked spikes are significantly suppressed or delayed, as shown for several trials in control (black) or with paired optical stimulation (blue). **(C)** Summary data showing cholinergic-mediated suppression of spiking suppression (n = 10 cells).

